# The spatiotemporal effects of seasonal migration on passerine phylogenetic community structure

**DOI:** 10.64898/2026.08.18.745533

**Authors:** Matthew D. Hack, Benjamin M. Winger

## Abstract

1. Seasonal migration in birds involves a substantial spatial redistribution of avian biodiversity each year and drives seasonal changes in community composition. Migrants experience different combinations of species interactions over space and time, generating regular disassembly and reassembly of bird communities throughout their annual cycles. However, the effects of seasonal migration on phylogenetic community structure remain poorly understood.
2. We assess spatiotemporal variation in phylogenetic community structure of North American passerines to test how seasonal migration restructures the evolutionary relatedness and dominant assembly mechanisms in bird communities throughout the annual cycle. Using distributional projections, we calculated metrics describing the phylogenetic dispersion of passerine communities each week of the year. We then tested the relationship between seasonal turnover in community phylogenetic dispersion and seasonal variation in species richness and proportion of migratory species.
3. Seasonal migration, by changing spatial patterns of avian diversity, simultaneously drives a complex continental redistribution of phylogenetic community structure. We find evidence of taxonomic scale dependency to our results, wherein throughout North America, the seasonal influx of migrant passerines yields communities that are overall more phylogenetically clustered, yet also exhibit greater phylogenetic overdispersion at smaller taxonomic scales.
4. Seasonal shifts in phylogenetic dispersion, though complex, track changes in diversity, manifesting as fluctuations in phylogenetic dispersion between northern and southern regions as seasonal migrants move between these regions. Our findings reveal a dynamic continental landscape of phylogenetic community structure directed by the movements of seasonal migrants.

## Introduction

The geographic distributions of species are determined by a combination of evolutionary history, environmental conditions, and species interactions (Ricklefs, 1987; Sexton et al., 2009). To understand the ecology and evolution of species distributions, it is critical to consider the relative influence of abiotic and biotic factors that shape the makeup of communities (Wisz et al. 2013). Studies of community phylogenetic structure (Cavender-Bares et al., 2004; Webb et al., 2002) have yielded insights into the interplay between evolutionary relationships and community assembly (Graham et al., 2009), which collectively influence species co-occupancy and range-wide distributions (Gotelli et al., 2010). However, most research has examined communities under the assumption that species compositions and interactions are temporally static (Emerson & Gillespie, 2008). This approach cannot capture the effects of seasonal migration on community structure throughout the year, as the movements of migratory species shift local species compositions as migrants carry out their annual journeys (Klingbeil & Willig, 2016).

Among birds in particular, the widespread prevalence of seasonal migration poses a unique challenge for investigating the factors underpinning avian community composition (Jarzyna & Stagge, 2023). The annual movements of seasonally migratory birds, which travel between separate breeding and non-breeding distributions, bring about profound spatial redistributions of avian biodiversity multiple times per year (Somveille et al., 2013). The estimated 19% of extant bird species that undergo seasonal migration (Kirby et al., 2008) contribute to substantial and predictable seasonal turnover in avian community composition and species richness (Hurlbert & Haskell, 2003; Keyser et al., 2024). These effects are particularly pronounced in the highly seasonal environments at higher latitudes and elevations (Newton & Dale, 1996), which lose most of their diversity as migrants depart during colder months (Somveille et al. 2013). This drastic turnover in community composition and richness could rearrange the forces driving the assembly of these communities. Though it is well-understood that the movements of migrants yield seasonal rearrangements of avian biodiversity and alter community composition (Ng et al., 2022), it remains less clear how seasonal pulses of migrants reorganize the phylogenetic structure of communities. Do migration-driven changes in community composition cause parallel redistributions of phylogenetic community structure patterns? Or do bird communities shift in species richness and membership while maintaining similar phylogenetic structure, suggesting consistency in their assembly processes through time and space (Fig. 1)? Here, we characterized the phylogenetic community structure of North American passerine bird assemblages throughout the annual cycle to assess the influence of seasonal migration on passerine community assembly.

**Figure 1:**
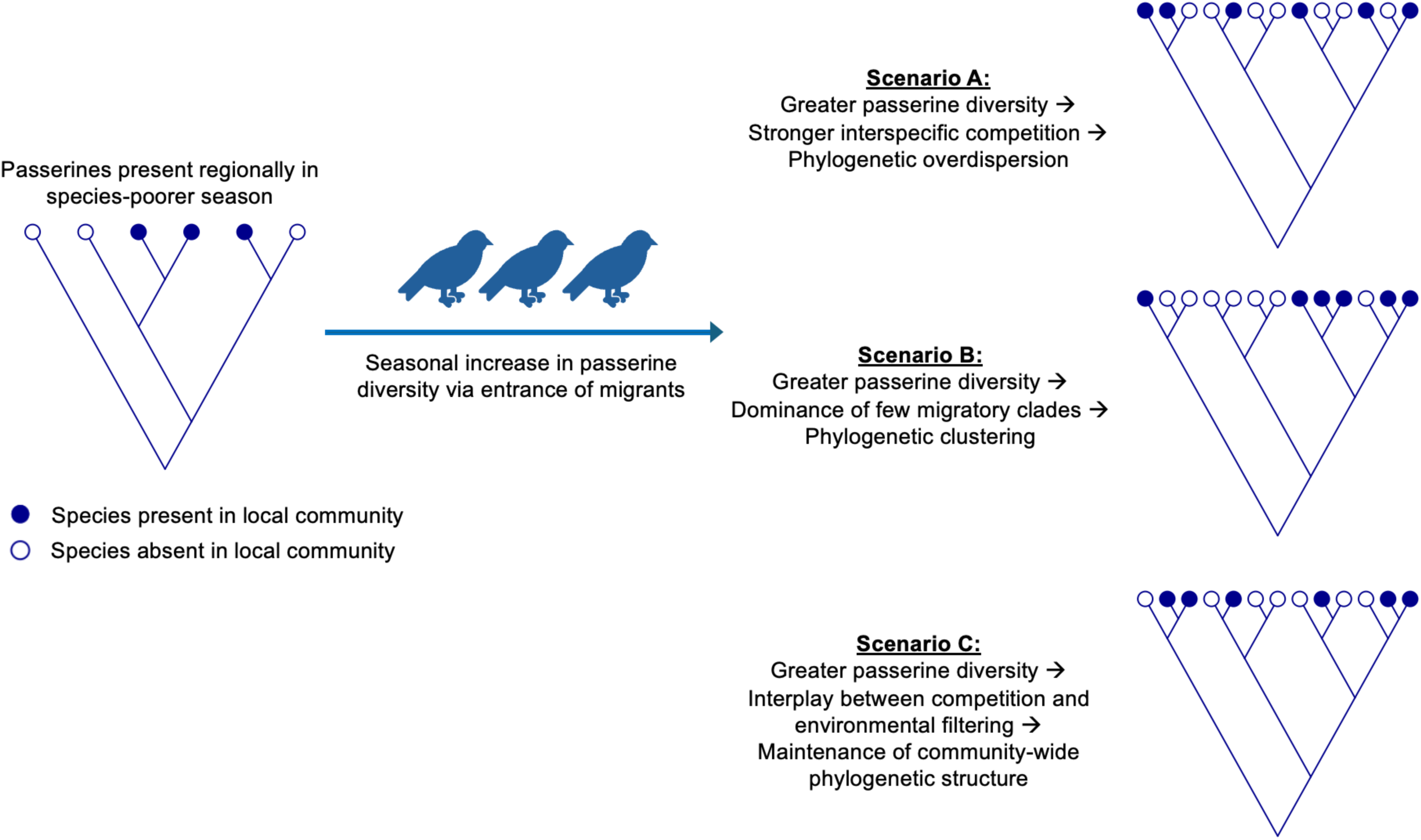
Conceptual model depicting possible effects of an influx of migratory birds on community phylogenetic structure. A seasonal increase in passerine diversity may or may not alter the relative strengths of assembly forces in structuring community occupancy. If the arrival of additional diversity is associated with increased interspecific competition, the community may become more phylogenetically overdispersed (Scenario A); alternately, the seasonal dominance of a small number of diverse clades may bring about a more phylogenetically clustered community (Scenario B). However, seasonal changes to community composition and diversity need not necessarily modify local phylogenetic community structure if assembly processes nonetheless remain consistent or counteract one another (Scenario C).

Community assembly is the outcome of several factors operating at multiple spatial and temporal scales, including historical biogeography, climate, species-specific habitat preferences, and interspecific interactions (Weeks et al., 2016; Weiher & Keddy, 2001). Though each of these factors influences the distributions and co-occurrences of species, their relative strength in shaping community assembly varies (Simberloff et al., 1999). The framework of phylogenetic community ecology has helped identify drivers of community assembly by using the phylogenetic structure of communities and trait evolution patterns to test mechanisms underlying community assembly patterns (Cavender-Bares et al. 2004; Cavender-Bares et al. 2009; Webb et al. 2002).

Most quantifications of phylogenetic community structure compare the degree of phylogenetic relatedness among co-occurring assemblage members against that of randomly simulated communities derived from a regional species pool (Carstensen et al., 2013). Communities that demonstrate more phylogenetic distance between sympatric species than null expectation are considered phylogenetically overdispersed; meanwhile, phylogenetically clustered communities show less phylogenetic distance than expected. Phylogenetic clustering may indicate a predominant role of environmental filtering, whereby climatic conditions and evolutionary history determine species’ distributions according to their shared, phylogenetically conserved traits (Cavender-Bares et al., 2004). By contrast, phylogenetic overdispersion may reflect interspecific competition among closely related species, potentially implicating biotic constraints in preventing species from occupying the entirety of their climatic niche (Hargreaves et al., 2014).

Assembly mechanisms may act unevenly at different taxonomic scales. For example, competitive exclusion could structure community composition primarily among close relatives (González-Caro et al., 2012). A nuanced interpretation of community dynamics thus calls for a comparison between multiple phylogenetic dispersion metrics. Two commonly used metrics are net relatedness index (NRI), which captures evolutionary distance across all combinations of co-occurring taxa, and nearest taxon index (NTI), which measures evolutionary distance specifically between closest co-occurring relatives (Webb, 2000). While NRI and NTI are often correlated, their differences can elucidate community assembly mechanisms by specifying the taxonomic scopes at which environmental filtering and competitive exclusion predominate.

Migratory birds provide an opportunity to test the more general effects of phenologically varying species richness on phylogenetic community dynamics, while holding the geographic location of the community itself constant. As seasonal migrants travel, they may experience divergent patterns in phylogenetic structure between their two stationary seasonal communities (breeding and non-breeding seasons—hereafter “stationary seasons”); likewise, their entrance and exit from these communities may involve different assembly mechanisms. For example, a migrant-driven increase in species richness may involve increased interspecific competition and yield a seasonal turn towards phylogenetic overdispersion (Powell et al., 2021). Yet seasonal migration can exhibit phylogenetic signal, as closely related species often demonstrate similar migratory behavior (Winger et al., 2012); temporal variation in species richness could thus be largely driven by several diverse families with many migrant species (Rabenold, 1993), thereby potentially yielding phylogenetic clustering.

The biogeographic gradients of North American passerine diversity affords an opportunity to directly compare migrant-dominated assemblages that vary seasonally. Critically, within this region, passerine species richness peaks in northern regions during the breeding season and in southern regions during the non-breeding season, with a previously documented transition zone between these regions around 35°N (Somveille et al., 2013). This transition reflects a steep gradient in temperature seasonality and facilitates a comparison between regions with migrant concentrations that peak in the summer or the winter. The North American passerine assemblages thus make an ideal system for a rigorous quantification of community phylogenetic structure throughout the annual cycle needed to better understand the phylogenetic dynamics of seasonal assemblages.

We aimed to describe and quantify the influence of seasonal migration on the phylogenetic structure of assemblages throughout the annual cycle. We predicted differences in community phylogenetic structure between the stationary seasons to covary predominantly with seasonal changes in species richness, as changes to community composition can be understood as diversity “pulses” of migrants entering and exiting. More specifically, we predicted local communities will become more phylogenetically overdispersed during whichever stationary season contains more migratory species (Somveille et al., 2019) due to space partitioning between close taxonomic and ecological relatives. We thus expect greater phylogenetic overdispersion during the summer at higher latitudes, and during the winter at lower latitudes. Alternately, if migration is phylogenetically conserved (Alerstam et al., 2003; Winger et al., 2019), migrant-dominated assemblages may become inundated with close relatives and therefore become more phylogenetically clustered. In describing the spatiotemporal variation in passerine phylogenetic community structure, our framework helps to clarify the role of seasonal migration in reorganizing passerine community dynamics.

## Materials and Methods

### Study system selection

We restricted our study to temperate North America, allowing us to test the interplay between seasonal migration and community structure in a biogeographic region whose avian assemblage is strongly influenced by seasonal migration (Somveille et al., 2013). The divide between the Nearctic and the Neotropics represents a biogeographic division between avian systems dominated by seasonal migrants and residents (Cox, 1985), allowing us to focus on a system where the effects of migration-driven assemblage turnover are not overwhelmed by year-round residents (Bell, 2000). We chose to exclude the Neotropics as high-quality distributional projections (described below) are not yet available for their complete passerine assemblage.

We carried out phylogenetic community analyses on the passerine assemblages of the continental United States and Canada, comprising 297 species (227 migratory and 70 non-migratory species), representing all passerines that occur regularly in the terrestrial habitats of this region, excepting eight species for which robust distributional projections were unavailable. We selected passerines due to their diversity of seasonally migratory species, which disproportionately comprise the set of terrestrial migrants that contribute to seasonal community turnover (Rabenold, 1993), and represent the vast majority of plausibly interacting avifauna competing for overlapping resources. Although passerines interact ecologically with non-passerine birds in various ways, the inclusion of nonpasserines would also include non-interacting taxa that would bias findings towards phylogenetic clustering, along with predators at different trophic levels, so we restricted the study to passerines to improve interpretability of results. We used species distributional models produced by the eBird Status and Trends Data Products (“eBirdst”), which applies citizen science-driven bird observations and satellite imagery to infer species distributions at fine-grained precision (Fink et al., 2023). These models, which project occurrence and abundance throughout species’ ranges during each week of the year, are available for most North American passerines, excluding a small number of vagrant or highly range-restricted species.

### Generating weekly communities

We used the R package ‘ebirdst’ (Fink et al., 2022) to download species-specific mean relative abundance distributional model rasters from the ‘ebirdst’ 2021 data version (Fig. 2). We modeled local assemblages by creating weekly distributional rasters for each species and aggregating rasters from native 2.96×2.96 km resolution into 5×5 km hexagonal grids (following Gotelli et al. 2010 as an appropriate passerine community size estimate enabling the detection of interspecific competition; Fig. 2). Each hexagon represents one local community of passerines during one week of the year. We used these grids to generate a weekly continent-wide matrix of passerine presence in each simulated community. Approximately 1.1-1.3 million local communities were generated per week, depending on the available spatial extent of weekly eBirdst models, and trimmed to the extent modeled for all included species. We calculated the proportion of community diversity comprised of seasonal migrants (hereafter “migratory proportion”), by classifying each species based on migratory status (a binary migratory-vs.-non-migratory classification) following Pegan & Winger (2020). We classified any species containing migratory populations within North America as “migratory”. Though many North American migrants are partially migratory (Torstenson et al., 2024), we classified these species as “migratory” as their movements contribute to seasonal turnover, and because distributional models showing year-round presence cannot distinguish between sedentary populations, partial migration, and a complete replacement of individuals.

**Figure 2:**
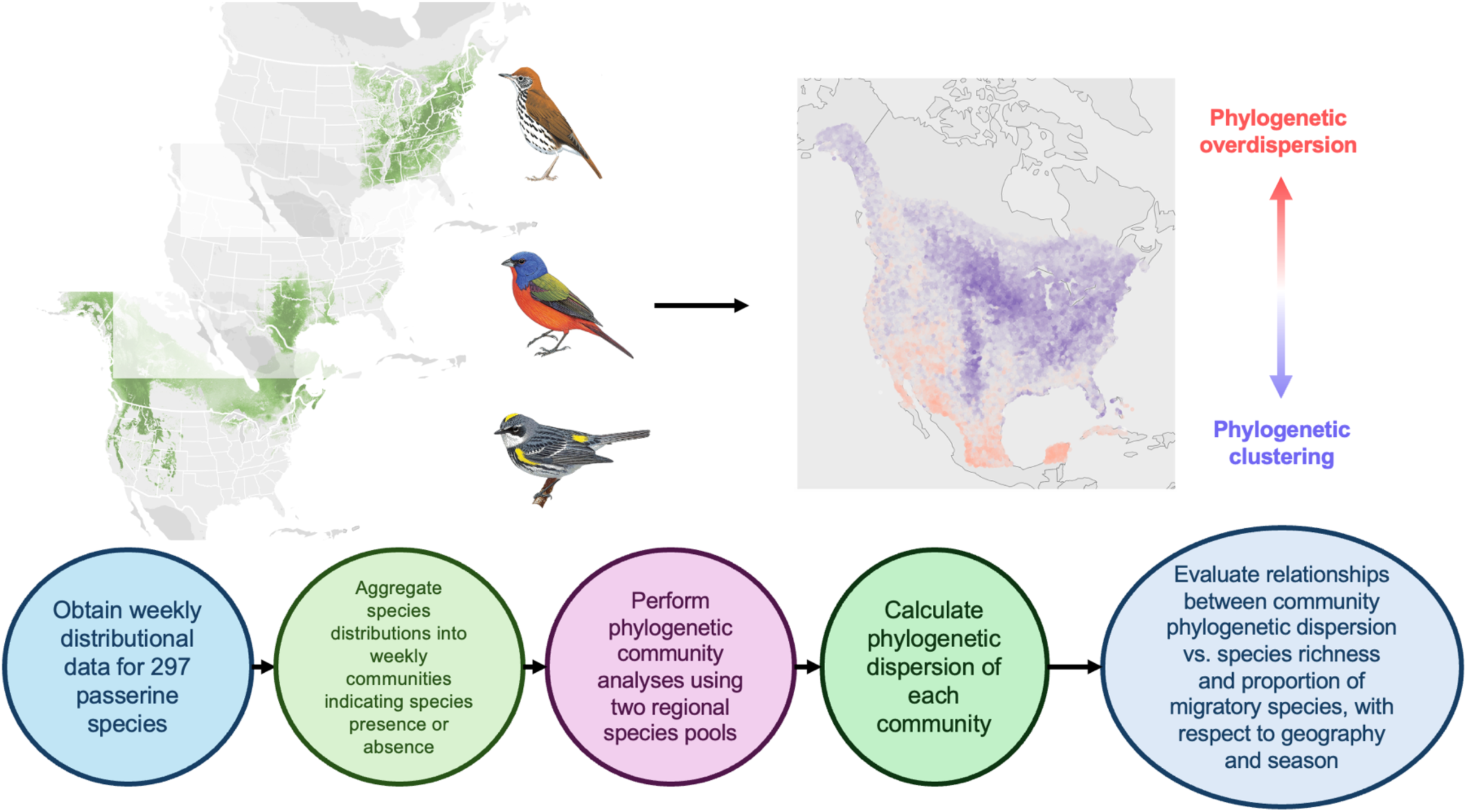
Conceptual flowchart outlining methodological framework. We obtained weekly species-specific distributional data and generated weekly passerine communities describing species presence and absence at the 5×5 km scale. We performed phylogenetic community analyses on each weekly community to calculate their phylogenetic dispersions using the metrics net relatedness index and nearest taxon index, using two regional species pools at different geographic extents. These metrics quantified the relationships between community phylogenetic patterns and seasonal migration over space and time. Bird illustrations courtesy of Cornell Lab of Ornithology, illustrators Brian Small, David Quinn, and Tim Worfolk from Birds of the World (Billerman et al., 2020).

### Regional species pool selection

For all community phylogenetics analyses, we compared weekly local passerine assemblages against simulated assemblages drawn from a regional species pool. The choice of the regional species pool can greatly affect quantifications of phylogenetic community structure (Lessard, Belmaker, et al., 2012). The filtering process between the species present at a continental and local scale involves historical processes of diversification and dispersal on an evolutionary time scale, whereas species pools capturing smaller regions preferentially infer local ecological processes (Lessard, Borregaard, et al., 2012; Zobel, 1997). To address the possibility of macroevolutionary and ecological processes indicating conflicting patterns in community phylogenetic structure, we calculated all metrics using regional species pools at two spatial extents (Fig. 2), both based on species presence inferred through ‘ebirdst’ models: a weekly continental pool comprising each species present during the given week at any point continentally; and ecoregion-level pools, using Bird Conservation Regions (BCRs) identified by the North American Bird Conservation Initiative (North American Bird Conservation Initiative, 2020). BCRs are ecoregions delineated by patterns in bird species distributional boundaries and habitat breaks, such that one BCR characterizes relatively homogenous bird communities and habitat (Sauer et al., 2003), thereby better capturing the role of ecological processes in filtering to local communities. We used BCRs 1-37, which encompass our study region. Each weekly ecoregional species pool contains all species present anywhere within the given BCR and week.

### Phylogenetic community metrics

We performed all phylogenetic analyses using a published comprehensive species-level phylogeny of New World Passeriformes, which contains over 2,500 species (Jetz et al., 2012). We generated a 50% majority-rule consensus tree via the SumTrees function in DendroPy (Sukumaran & Holder, 2010), following Rubolini et al., 2015. We used the R package ‘ape’ (Paradis & Schliep, 2019) to prune the complete Passeriformes tree to the set of species present in each regional community. All community phylogenetic analyses were performed using the R package ‘picante’ (Kembel et al., 2010). Phylogenetic dispersion metrics can consider each pairwise phylogenetic distance equally, or preferentially weight patterns concentrated at the level of the family or genus (the “tips” of the phylogeny; Kembel & Hubbell 2006; Kraft et al. 2007); these values are often correlated but may diverge depending on the specific pattern of phylogenetic dispersion (Vamosi et al., 2009). As described above, the two most common metrics for measuring phylogenetic dispersion are NRI, which measures the mean evolutionary distance between combinations of sympatric community members and better detects environmental filtering, and NTI, which captures the mean evolutionary distance between each combination of a species and its closest co-occurring relative and better detects competitive exclusion (Freilich & Connolly, 2015; Webb, 2000). As either filtering or competition could plausibly or jointly shape assembly, and as competition strength may vary with respect to taxonomic scale (Burns & Strauss, 2011), a comparison between both metrics enables a more precise analysis of community dynamics.

To calculate dispersion metrics, we created phylogenetic variance-covariance matrices for each regional community to calculate the evolutionary distance between each combination of taxa. We randomly subsampled 25,000 communities per week due to computational constraints. We used variance-covariance matrices to estimate community-level mean phylogenetic distance (MPD), along with mean nearest taxon distance (MNTD), which measures mean evolutionary distance between closest co-occurring relatives. MPD resolves phylogeny-wide patterns of phylogenetic community structure, while MNTD is more sensitive to clade-specific patterns concentrated at the terminal branch structure (Kembel & Hubbell, 2006; Kraft et al., 2007). We corroborated this assumption by calculating the mean phylogenetic distance of each pairwise species combination captured by each metric during both stationary seasons (measured as the first week of January and the last week of June, when North American passerine assemblages are relatively static; Fig. S1). We compared community MPD and MNTD against 100 simulated permutations of the regional species pool (Kembel, 2009) and calculated NRI and NTI as the standardized effect size of MPD and MNTD, respectively, against the null distribution multiplied by-1 (Webb, 2000; Fig. 2). For both NRI and NTI, positive and negative values represent phylogenetic clustering and overdispersion, respectively.

### Statistical modeling and seasonality

We assessed the influence of passerine species richness and migratory species proportion on community-level seasonal variation in NRI and NTI (hereafter ΔNRI and ΔNTI; Figs. 2 & 3). As we hypothesized that the seasonal influx of migrants leads to increased competition, we predicted an association between community species richness (representing migrant-driven changes to community composition) and phylogenetic overdispersion (that is, seasonal increase in richness yields decreased ΔNRI and ΔNTI values). We modeled ΔNRI and ΔNTI as functions of seasonal turnover in species richness and migratory proportion between stationary seasons. Species richness and migratory proportion are correlated but not substitutable; the species present during the less diverse season may represent either residents, or distinct migratory species that turn over seasonally. The variance inflation factors of the GLMMs containing turnover in both species richness and migratory proportion ranged from 1.82 to 2.04, suggesting sufficiently low collinearity to avoid unstable estimates when jointly considering these predictors.

**Figure 3:**
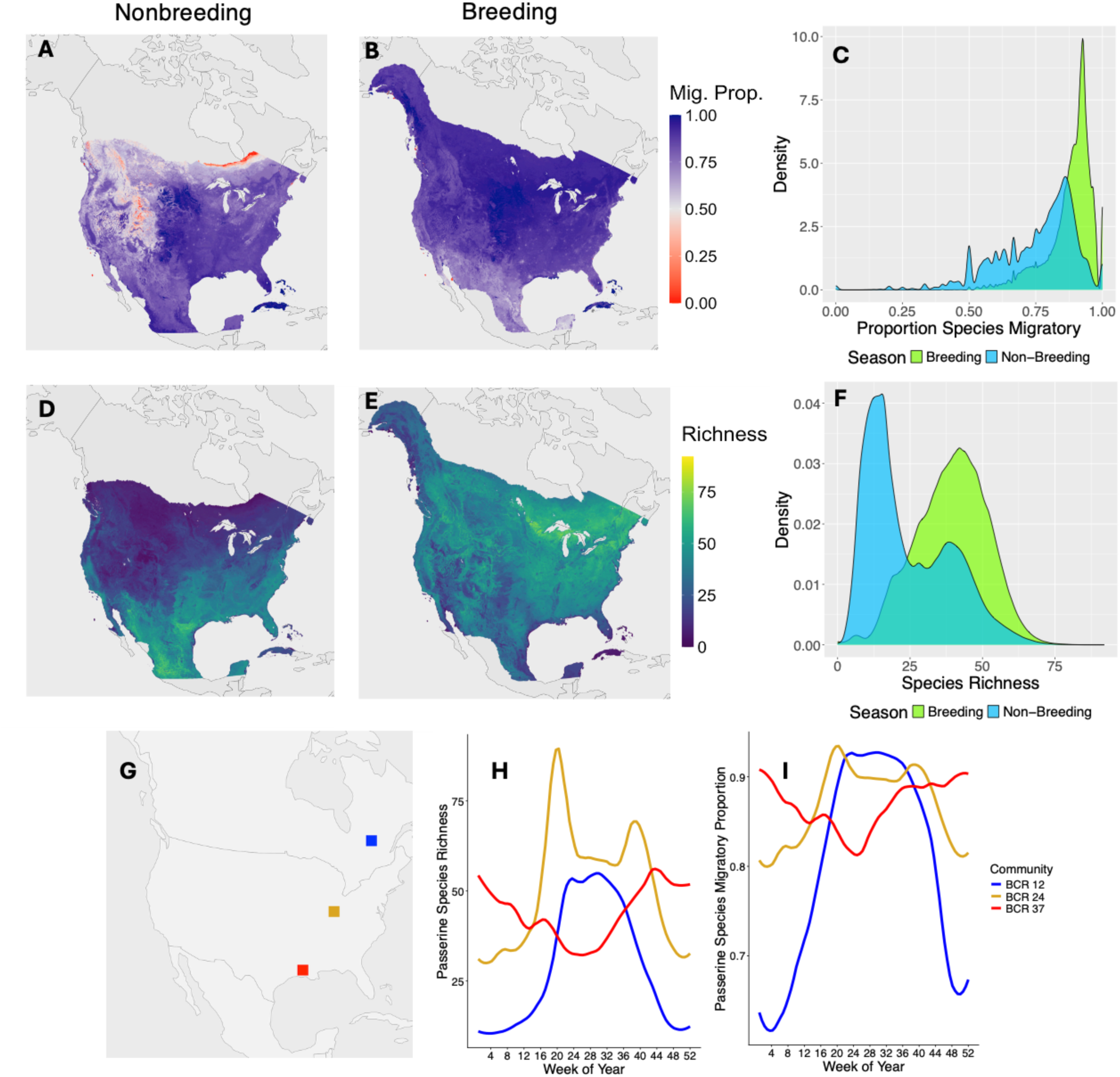
Spatiotemporal variation in patterns of migratory species diversity in North American passerine communities. (A, B) describe the community’s proportion of species that are migratory during the first week of January (A), representing the Northern Hemisphere nonbreeding season, and the last week of June (B), representing the breeding season, across the seasonal spatial extent available in eBird Status and Trends. Though most communities are composed predominantly of migratory species throughout the year, this proportion generally increases during the breeding season (C), particularly at higher latitudes and elevations. (D, E) represent patterns in species richness during the nonbreeding (D) and breeding (E) seasons. Overall, species richness increases during the breeding season (F), but the variation lies along a latitudinal gradient, with large diversity gains at higher latitudes and decreases at some southern latitudes. The temporal variation in community composition in three geographically disjunct communities (G) is shown in (H, I). (H) represents variation in species richness throughout the annual cycle; (I) represents seasonal variation in proportion of the community comprised of migrants.

There exists significant spatial autocorrelation in ΔNRI and ΔNTI—Moran’s I ranged from 0.13 to 0.32 (p<2.2e-16) across the dispersion metric-species pool combinations—thus necessitating spatial models. To account for spatial autocorrelation among communities, we tested relationships via spatial generalized linear mixed models (GLMMs), which include a spatial random field. We constructed GLMMs using the R package ‘sdmTMB’ (Anderson et al., 2025). To test the individual and joint influences of species richness and migratory proportion on ΔNRI and ΔNTI, we fit three GLMMs per dispersion index: separate models using either species richness or migratory proportion as predictors, and a model including both predictor variables and an interaction term.

Seasonality plays a central role in shaping the geography of migration (Winger et al., 2019), and geographic variation in temperature seasonality is tightly linked to observed gradients in species richness and migratory behavior. We thus investigated the influence of temperature seasonality on the extent of seasonal turnover in phylogenetic dispersion. We used the BioClim database (Busby, 1991) to extract two bioclimatic variables capturing local seasonality: BIO4 (temperature seasonality) and BIO7 (annual temperature range). We tested the influence of seasonality on absolute value of ΔNRI and ΔNTI by fitting spatial GLMMs using BIO4 and BIO7 as predictors.

### Data availability statement

All analyses were conducted using R Statistical Software (R Core Team, 2022). The data associated with this manuscript are available for peer review in Dryad (https://doi.org/10.5061/dryad.gf1vhhn2g) (reviewer link: http://datadryad.org/share/5wcPZKSkaDs3v0fAFQ4zbXAkqSv_LNHBmQStt_IyHKI). The code associated with this manuscript is available for peer review in Zenodo (DOI 10.5281/zenodo.17094330) (reviewer link: https://zenodo.org/records/17094330?preview=1&token=eyJhbGciOiJIUzUxMiJ9.eyJpZCI6IjRlMDcyMzFmLTVjYzMtNGRmNy1iZWEzLTAzNTdmNzZkNzY5MiIsImRhdGEiOnt9LCJyYW5kb20iOiIzNDliZWQ1Yzk3Y2M2MzVjNTI3MDk5OGFjYTRlMmU1ZSJ9.Q0VgOLftMOqRWPKhzUd7OBA8L8M7KTrYsLIPUErTuKuzcXwskzqnwjjLMusVp2gU021H47vsY_VmiWlwsuS0Eg). Data and code will be made publicly available upon acceptance.

## Results

### Continental patterns in passerine community diversity

Median passerine species richness was 34 species (SD=16.18); communities ranged from 1 to 122 species present (Fig. 3). Migratory proportion varied from 0 to 1, with a median of 0.86 (SD=0.12), ranging from 0 to 116 migratory species (median = 29, SD=15.57). Species richness and migratory proportion are positively correlated (Pearson’s correlation = 0.56, p<2.2e-16). Overall, passerine species richness increased during the Northern Hemisphere summer; from January to June, mean species richness increased from 23.34 (SD=13.85) to 40.08 species (SD=11.62) (Figs. 3, 4C-D). Summer communities saw a parallel increase in migratory proportion (summer mean = 0.88, SD=0.08; winter mean = 0.75, SD=0.15). However, seasonal changes in passerine diversity were neither uniform in directionality nor magnitude. Higher latitudes experienced strong increases in species richness and migratory proportion during the summer, while lower latitudes demonstrated diminished or reversed patterns (Fig. 4C-D).

**Figure 4:**
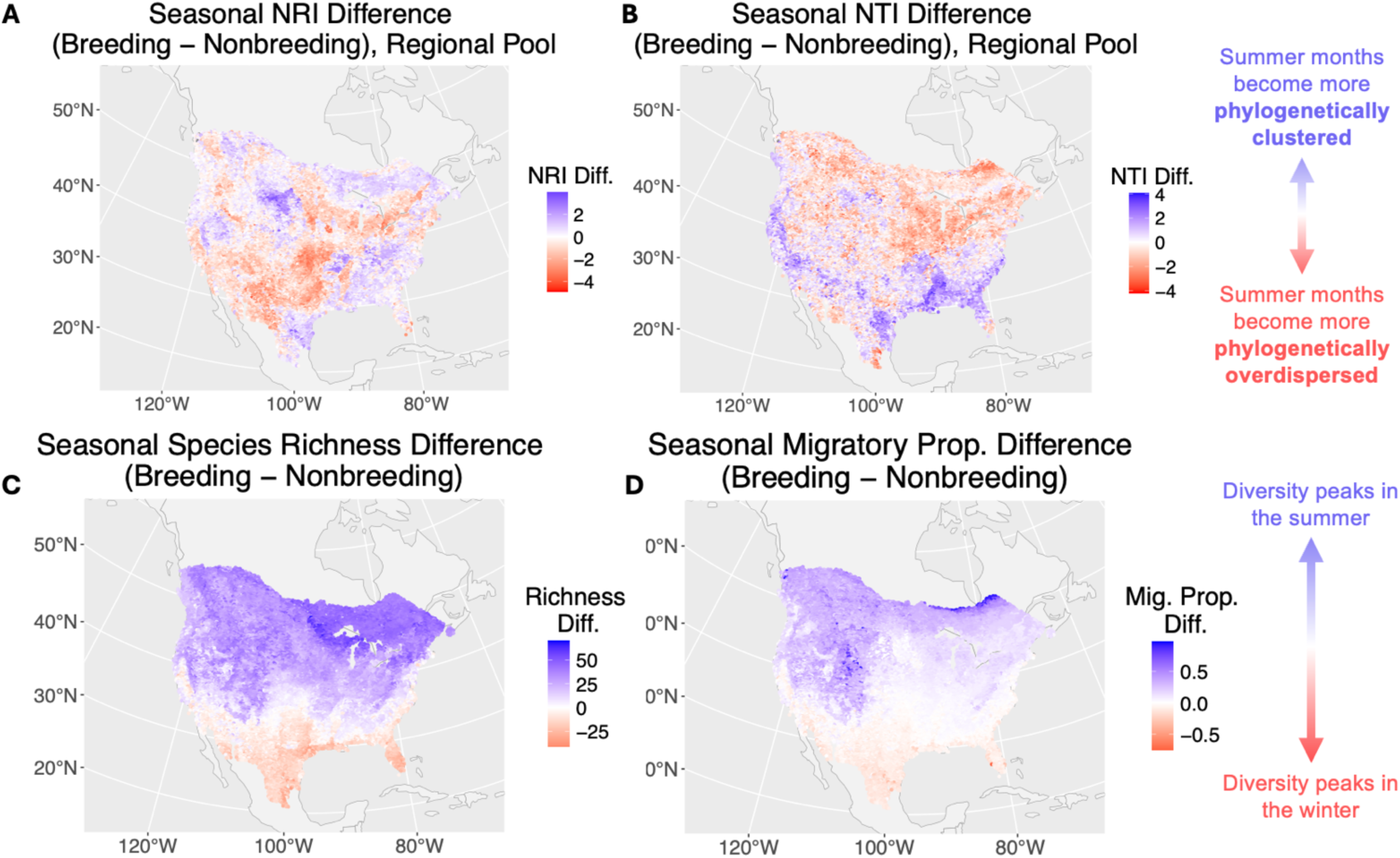
Maps representing community seasonal differences in net relatedness index (NRI; A); nearest taxon index (NTI; B); species richness (C); and migratory proportion (D). Results show phylogenetic dispersion-community composition relationships using the ecoregional species pool, and results using a continental species pool are shown in Fig. S4.. Seasonal difference values represent breeding season values (using data from the last week of June) subtracted by non-breeding season values (first week of January). For both indices, blue values represent greater phylogenetic clustering during summer as compared to winter; red values depict greater phylogenetic overdispersion during summer. Blue and red values also represent higher species richness (C) and migratory proportion (D) during the breeding and non-breeding seasons, respectively.

### Patterns in phylogenetic community structure variation

Results from using ecoregional (BCR-based) species pools are presented here and results from the continental species pools are presented in the SI; results between the two species pools were consistently qualitatively similar (Tables 1, S1) Under ecoregional models, NRI averaged - 0.27 (SD=1.04), indicating a slight tendency towards phylogenetic overdispersion on average, while NTI averaged 0.32 (i.e., slight phylogenetic clustering; SD=0.97). NRI and NTI exceeded 0 in 38.9% and 62.6% of communities, respectively (Fig. S2). Phylogenetic distances captured by NRI and NTI remained consistent seasonally (Fig. S1); the mean pairwise distance (in estimated Mya since divergence) among species combinations captured in NRI was 80.11 in January and 86.26 in June; for NTI, 22.60 and 19.66, respectively. NTI thus captures community phylogenetic patterns between species at, on average, approximately a quarter of the phylogenetic distance as does NRI.

### Migrant-driven community turnover influences phylogenetic community structure

Seasonal changes in community composition driven by seasonal migration yield predictable shifts in phylogenetic community structure. A seasonal increase in either species richness or migratory proportion was associated with a parallel seasonal increase in NRI, but a seasonal decrease in NTI (Table 1; Figs. 5, S5). The directionality of these relationships was consistent when separately modeling species richness or migratory proportion, though the global ΔNTI model incorporating both predictors flipped migratory proportion signs, likely due to the moderate correlation between the predictors. The effect sizes of changes in migratory species richness on phylogenetic dispersion (Table 1) can be illustrated by comparing northern versus southern communities. For example, a change in species richness turnover between a southern BCR (e.g., BCR 37, the Gulf Coastal Prairie along the Gulf Coast, whose communities lose an average of 19 species in the summer) and a northern BCR (e.g., BCR 12, the Boreal Hardwood Transition in the upper Great Lakes, which gains 45 species in summer) is associated with a predicted increase of 0.788 in ΔNRI (from-0.722 to 0.066), and a predicted decrease of 1.344 in ΔNTI (from 0.642 to-0.702). These values are slightly greater than one standard deviation of the total continental variation in NTI (SD = 0.959 in summer and 0.967 in winter), and slightly less than one standard deviation of NRI (SD = 0.863 in summer and 0.957 in winter; Fig. 4). An analogous change in migratory proportion turnover (-0.087 in BCR 37 to 0.342 in BCR 12) predicts similar results (ΔNRI increases from-0.612 to 0.121; ΔNTI decreases from 0.192 to-0.404). These results indicate that shifts in passerine migratory diversity yield nontrivial shifts in community phylogenetic dispersion. We replicated the same models using the continental species pool; predictor directionality was unchanged across species pools and coefficient magnitude was comparable, indicating model spatial grain did not alter our conclusions regarding any key relationships (Tables 1, S1; Figs. 5, S5).

**Figure 5:**
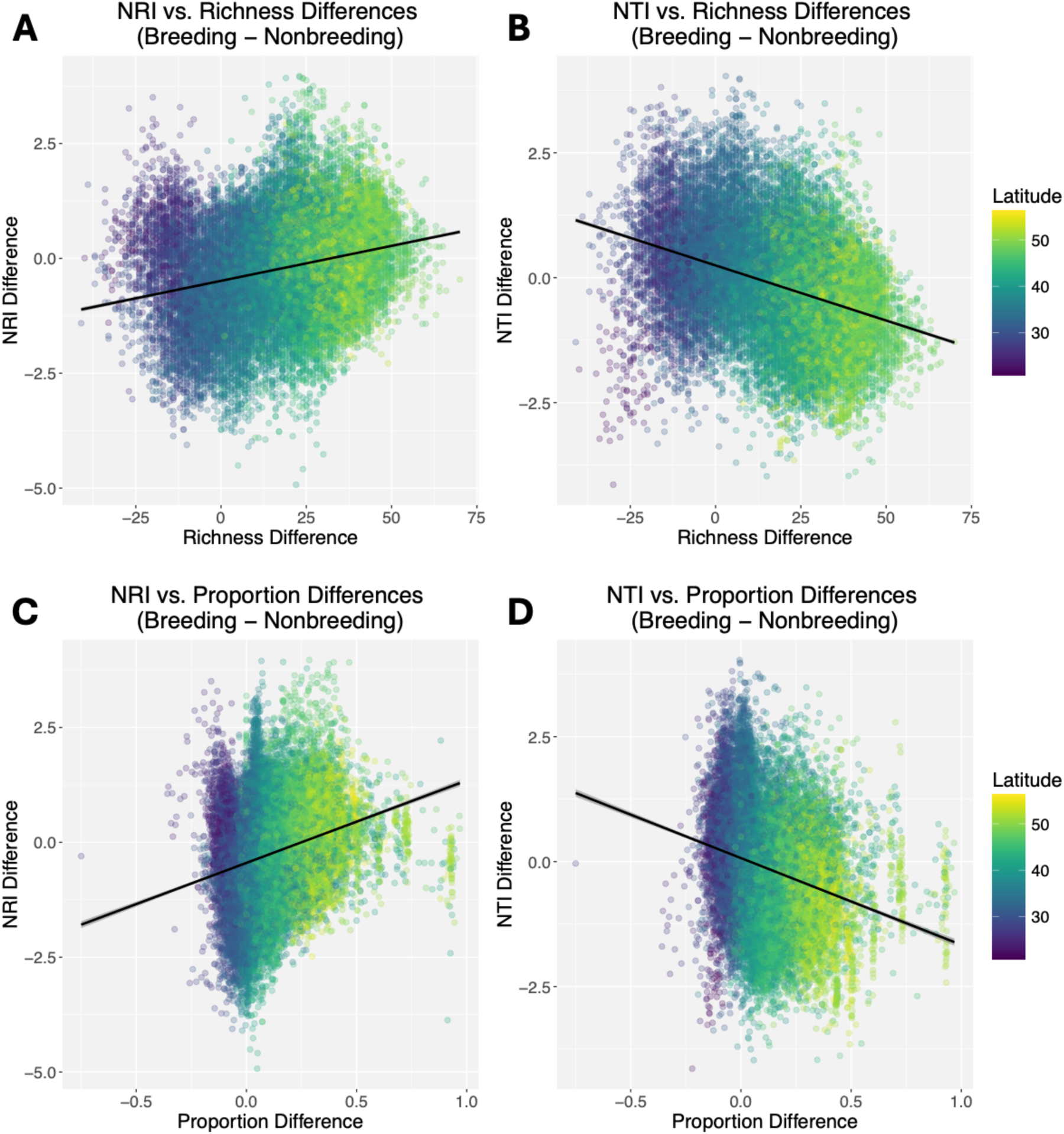
Scatterplots depicting relationships between seasonal differences in community dispersion, and seasonal differences in species richness (A, B) and migratory proportion (C, D). Results show phylogenetic dispersion-community composition relationships using the ecoregional species pool, and results using a continental species pool are shown in Fig. S5. (A, C) and (B, D) represent NRI and NTI relationships, respectively. Seasonal differences represent breeding season values (calculated as the last week of June) subtracted by non-breeding season values (first week of January). Spatial GLMMs predict that seasonal increases in richness and migratory proportion are associated with a seasonal increase in NRI (greater clustering) and decrease in NTI (greater overdispersion). Standard errors are very low such that error bars around trendlines are negligible.

**Table 1:** Results of spatial generalized linear mixed models quantifying drivers of variation in seasonal turnover in NTI and NRI, with predictor variables of seasonal turnover in species richness and migratory proportion (along with their interaction). These results are derived from the ecoregional species pool. Parameters whose 95% confidence intervals do not overlap with zero are bolded.

| ΔNTI | Term | Estimate | SE | 95% CI |
| --- | --- | --- | --- | --- |
| Richness | Intercept | 0.247 | 0.00962 | **[0.228, 0.266]** |
|  | Richness | -0.0212 | 0.000419 | **[-0.0221, -0.0204]** |
| Proportion | Intercept | 0.0710 | 0.00919 | **[0.0530, 0.0890]** |
|  | Proportion | -1.39 | 0.0541 | **[-1.50, -1.29]** |
| Global | Intercept | 0.251 | 0.00995 | **[0.231, 0.270]** |
|  | Richness | -0.0249 | 0.000565 | **[-0.0260, -0.0238]** |
|  | Proportion | 0.989 | 0.103 | **[0.786, 1.19]** |
|  | Richness x Proportion | -0.0120 | 0.00312 | **[-0.0181, -0.00584]** |

**Table 1:** Results of spatial generalized linear mixed models quantifying drivers of variation in seasonal turnover in NTI and NRI, with predictor variables of seasonal turnover in species richness and migratory proportion (along with their interaction).
| ΔNRI | Term | Estimate | SE | 95% CI |
| --- | --- | --- | --- | --- |
| Richness | Intercept | -0.490 | 0.0101 | **[-0.510, -0.471]** |
|  | Richness | 0.0124 | 0.000441 | **[0.0115, 0.0132]** |
| Proportion | Intercept | -0.463 | 0.0092 | **[-0.481, -0.445]** |
|  | Proportion | 1.71 | 0.0542 | **[1.61, 1.82]** |
| Global | Intercept | -0.529 | 0.0103 | **[-0.549, -0.509]** |
|  | Richness | 0.00524 | 0.000589 | **[0.00409, 0.00640]** |
|  | Proportion | 0.377 | 0.107 | **[0.167, 0.588]** |
|  | Richness x Proportion | 0.0354 | 0.00324 | **[0.0291, 0.0418]** |

Spatial GLMMs show temperature seasonality (BIO4) and annual temperature range (BIO7) to both have a significant positive association with absolute value of ΔNRI (Table S2). This suggests that more seasonal environments experience larger magnitudes of turnover in overall assemblage-wide relatedness than do less seasonal areas. Meanwhile, absolute value of ΔNTI is negatively associated with BIO7, and shows no relationship with BIO4, pointing to a less conclusive influence of climatic seasonality on variation in evolutionary relatedness among nearest relatives.

## Discussion

Our results show that seasonal migration, in driving spatiotemporal changes in songbird biodiversity, simultaneously gives rise to a parallel realignment of songbird community phylogenetic structure (Fig. S3). Throughout the annual cycle, the same geographic location is occupied by species assemblages with different degrees of phylogenetic relatedness; similarly, seasonal migrants themselves occupy assemblages with varying phylogenetic community structures in their breeding and non-breeding seasons. Thus, the seasonal movements of birds restructure community phylogenetic conditions for both migrants and residents.

The relationships between species richness, migratory proportion, and phylogenetic community structure illustrate the nature of migration-driven effects on passerine communities (Table 1; Fig. 5). A positive seasonal change in species richness or migratory proportion is generally associated with an increase in NRI, such that during the stationary season with greater migratory species richness, communities are more phylogenetically clustered when considering the entire passerine assemblage. This pattern is consistent with environmental filtering as an assembly process. However, when considering NTI, which more heavily weights phylogenetic relatedness among nearest co-occurring evolutionary neighbors, the more species-rich, migrant-dominated season is comparatively phylogenetically overdispersed. This pattern may reflect an increase in interspecific competition among closest relatives (Weber & Strauss, 2016).

We also find a strong geographic component to the complexity of seasonal variation in phylogenetic structure (Figs. 4 & 5). At higher latitudes, where passerine diversity peaks during the summer, the influx of migrants generates an assemblage characterized by closer relatedness at larger phylogenetic scales (higher NRI), but greater segregation of closest relatives into different communities (lower NTI). These richness-dispersion relationships are also observed at lower North American latitudes, but as their passerine diversity peaks during the winter, the seasonality of community phylogenetic dispersion flips—the non-breeding season experiences increased NRI and decreased NTI (Figs. 4 & 5). The geographic switch between these states closely aligns with the previously identified 35°N transition zone (Fig. 4; Somveille et al., 2013) marking the divide between regions experiencing peak diversity in opposite seasons. In essence, seasonal migrants transport the conditions of community-wide clustering and overdispersion among closest relatives as they move between their breeding and non-breeding grounds.

In tandem, these patterns suggest layered dynamics for how seasonal migration influences passerine community structure. Among North American passerines, migratory behavior is concentrated in certain diverse clades that disproportionately contribute to turnover in species richness—indeed, three diverse families (parulid warblers, passerellid sparrows, and tyrannid flycatchers) represent 50.2% of our dataset’s migratory species. Thus, because many migrant species from relatively few clades comprise an outsized proportion of a community’s species, the communities they enter become phylogenetically clustered. The positive relationship we find between seasonality and magnitude of NRI turnover (Table S2) shows that northern latitudes experience stronger reorganizations of phylogenetic structure, perhaps resulting from larger seasonal increases in available niche space (Keyser et al., 2024). These findings suggest that interspecific competition does not preclude coexistence among relatives within diverse migratory clades.

However, the opposite pattern in NTI suggests that within diverse clades, individual species tend to not co-occur with their closest evolutionary neighbors (Fig. S1), which may reflect competitive exclusion specifically structuring community assembly among closest relatives. The inconclusive relationships between seasonality and magnitude of NTI turnover (Table S2) suggest that phylogenetic structure at a fine scale more acutely reflects interactive processes than climatic seasonality. Under this framework, while environmental filtering outweighs competition in structuring co-occupancy, competition may nonetheless drive close phylogenetic neighbors into separate communities. This conclusion aligns with prior research in parulid warblers, which have high diversity in North America during summer but rarely co-occur directly with their regional sister taxa (Lovette & Hochachka, 2006).

The above interpretations consider the outcomes of migration between the stationary seasons (the northern summer and winter). During the spring and fall peaks of songbird transience, regional patterns in passerine community structure are much less evident (Fig. S3). Peak passerine migration is characterized by rapid assemblage turnover, perhaps at so brisk a pace that processes generally governing assembly are not evident. Songbird migratory stopover distributions are likely highly opportunistic and contingent upon habitat availability (Guo et al., 2024), topographic barriers, and weather conditions (Clipp et al., 2020; Van Doren & Horton, 2018). Passerines may also experience more positive interactions during migration, particularly in comparison to their breeding seasons (DeSimone et al., 2024). The contribution of these factors to the assembly of transient passerine assemblages calls for further research.

A key criticism leveled against community phylogenetics warns against inferring assembly mechanisms through phylogenetic structure alone (Davies, 2021; Lopez et al., 2016). Trait conservatism and niche conservatism are necessary assumptions that connect phylogenetic dispersion patterns to assembly drivers as postulated above; convergence and trait overdispersion can confound these results (Weber et al., 2017). Although seasonal migration can evolve rapidly (Rolland et al., 2014; Zink, 2002), it also exhibits phylogenetic signal in many of the passerine clades involved in this study (Gómez-Bahamón et al., 2020; Winger et al., 2012, 2014). While community phylogenetics cannot fully substitute for direct experimentation in demonstrating assembly mechanisms (Gerhold et al., 2015), our assumptions appropriately align with expectations with known phylogenetic patterns in seasonal migration. Additionally, our framework allows us to specifically identify *relative* shifts in phylogenetic dispersion patterns with respect to season, which circumvents several potential shortcomings in community phylogenetic analyses. Environmental filtering and interspecific competition jointly shape communities, (Lovette & Hochachka, 2006; Spasojevic & Suding, 2012), and their contributions can be obscured by an absolute classification of “overdispersion vs. clustering.” The temporal comparisons afforded by migration allow us to focus on the relative seasonal changes in community dispersion. Our confidence in these patterns is also bolstered by their robustness across multiple species pools. A potential pitfall of community phylogenetic metrics is their dependency on the spatial grain of the regional species pool (Lessard, Borregaard, et al., 2012), where smaller grains bias predictions towards phylogenetic overdispersion (Cardillo, 2011). However, our ecoregional models predicted comparatively overdispersed NTI but clustered NRI values (Fig. S2). More importantly, seasonal turnover in species richness and migratory proportion predicted parallel shifts in NRI and NTI across both spatial grains (Table 1A-B).

Our work highlights several paths forward to better understand the relationship between seasonal migration and community assembly. Most immediately, this study could benefit from the inclusion of lower tropical and subtropical latitudes, which contain far greater resident passerine diversity than does North America (Gómez et al., 2016), thus affording a larger gradient of species richness and migratory proportion. Though high-resolution Neotropical distributional models are increasingly available, they are not developed sufficiently across Neotropical passerine diversity (Feldman et al., 2021); the advancement of these models would enable the testing of our framework across a greater breadth of biodiversity and variation in migratory behaviors. Additionally, incorporating abundance data could move this analysis beyond community membership to capture changes in species dominance. Finally, future works could incorporate quantitative trait data, perhaps by generating community-level trait hypervolumes (Blonder, 2018) to test the contributions of migrants to niche packing or filling (Pigot et al., 2016), which could better elucidate the community effects of migration-driven species saturation.

Our study demonstrates that the combination of spatial variation in passerine diversity, and the migration-driven reorganization of their diversity patterns, contributes to a dynamic continental landscape of passerine phylogenetic community structure. As birds move geographically throughout their annual cycle, so too do the resulting phylogenetic structure patterns of the assemblages they enter and exit. Our findings also generate hypotheses regarding the effects of seasonal migration on community assembly mechanisms—in particular, field studies should test whether competitive exclusion between closely related taxa peaks during summers at higher latitudes and winters at lower latitudes. Ultimately, such work will contribute to a better understanding of the ecological effects exerted by seasonal migrants upon their communities.

## Notes

### Competing Interest Statement

The authors have declared no competing interest.

### Funding

- NSF DEB: 2146950

