## Supplemental Figures for "The spatiotemporal effects of seasonal migration on passerine phylogenetic community structure"

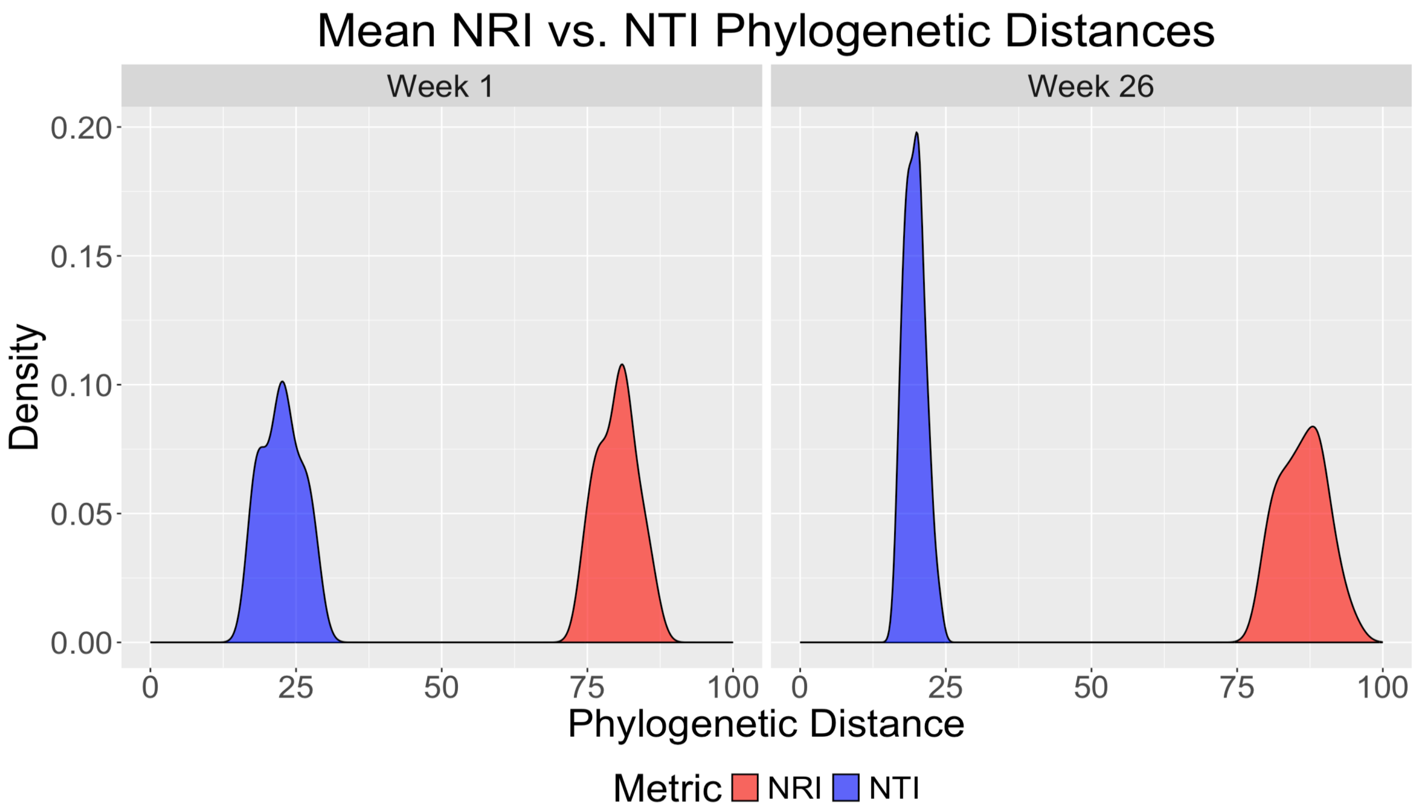


**Supplemental Figure S1:** Density plots depicting the distribution of BCR-level mean phylogenetic distances captured by NRI and NTI metrics during weeks 1 and 26. The phylogenetic distance values represent the mean branch length across all pairwise combination of species included in the relevant metric. The mean phylogenetic distances captured by NRI and NTI are consistent across the breeding and nonbreeding season, as does the ratio between the two metrics. These observations support the assumption that NTI more directly captures patterns within terminal branch structure.

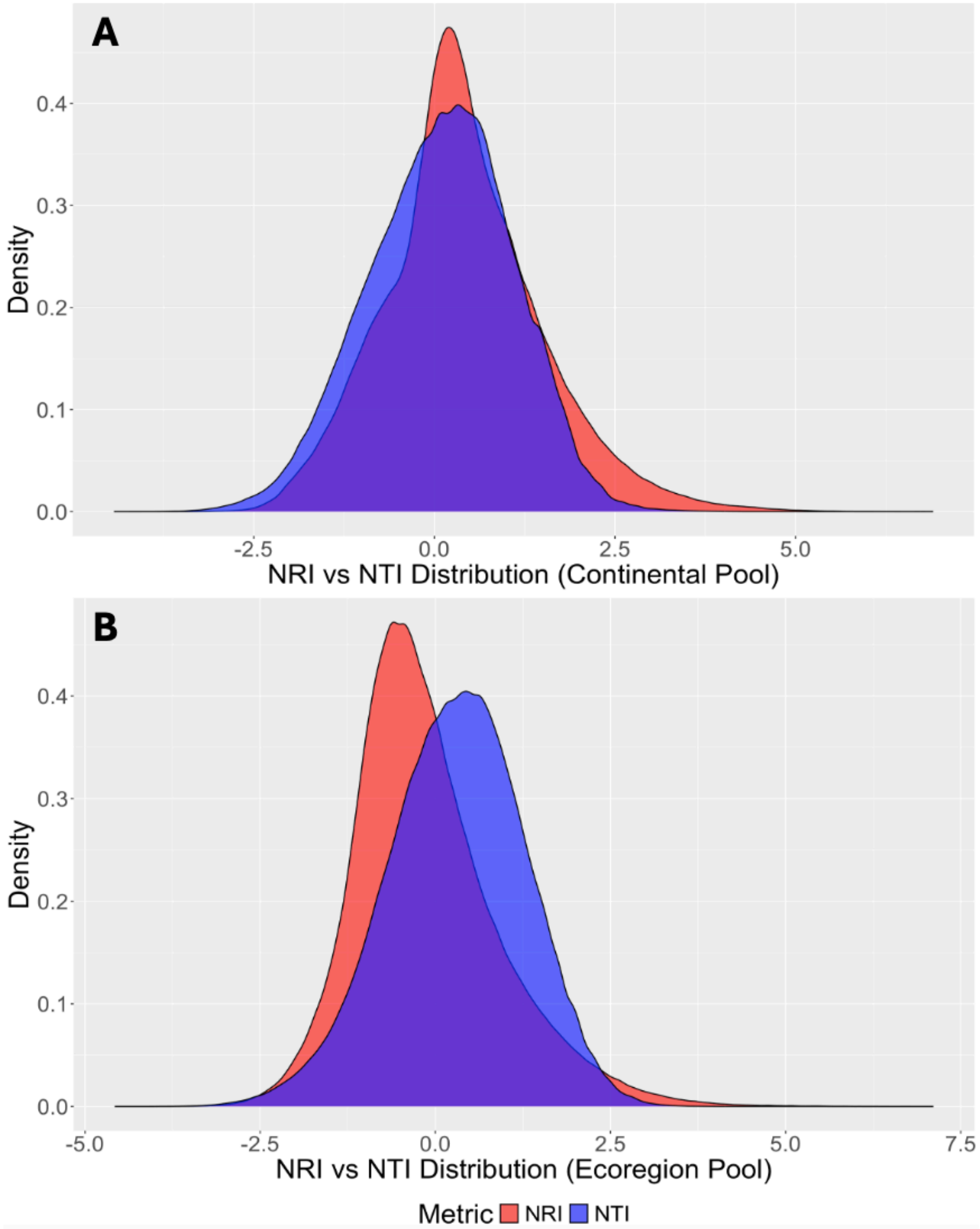
A
B
Density
0.4
0.3
0.2
0.1
0.0
-5.0
-2.5
0.0
2.5
5.0
7.5
NRI vs NTI Distribution (Continental Pool)
NRI vs NTI Distribution (Ecoregion Pool)
Metric
NRI
NTI

**Supplemental Figure S2:** Distributions of the phylogenetic dispersion metrics net relatedness index (NRI) and nearest taxon index (NTI) using the continental species pool (A) and the ecoregional species pool (B). At the continental pool, both NRI and NTI peak at values slightly over 0 (positive values represent phylogenetic clustering), though a greater proportion of NRI values than NTI values are positive. At the regional pool, NTI values peak slightly above 0, while NRI values peak slightly below 0, representing phylogenetic overdispersion.

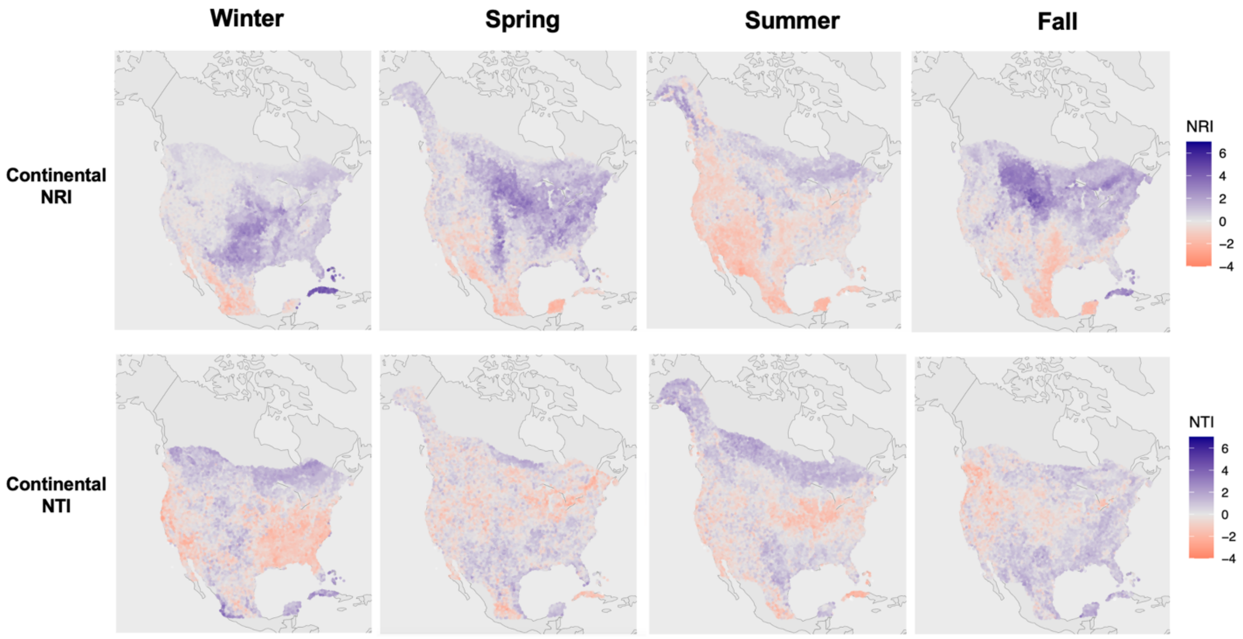

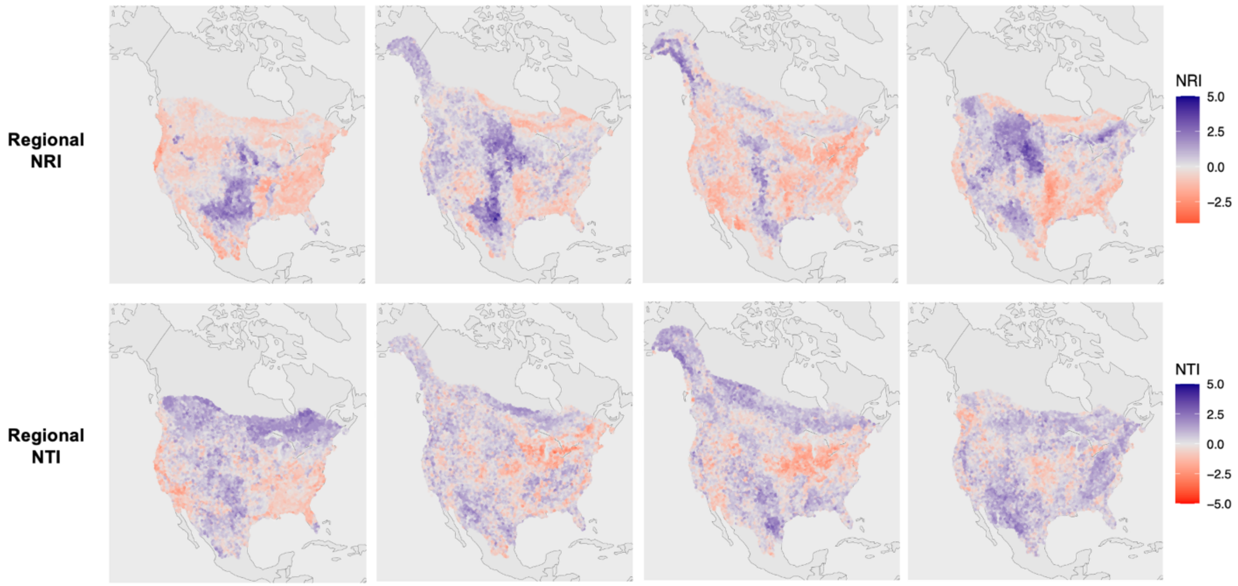


**Supplemental Figure S3:** Spatiotemporal variation in NRI and NTI values using the continental and ecoregional species pools. Columns represent phylogenetic dispersion values during (from left) winter, spring, summer, and fall, while rows represent phylogenetic dispersion using (from top) NRI and NTI values derived from the continental species pool, and NRI and NTI values derived from the ecoregional species pool. Communities colored in red are phylogenetically overdispersed, while communities in blue are phylogenetically clustered. In each model, migration-driven turnover in community diversity and composition brings about changes to continental patterns in phylogenetic dispersion, though the different models do not produce identical predictions.

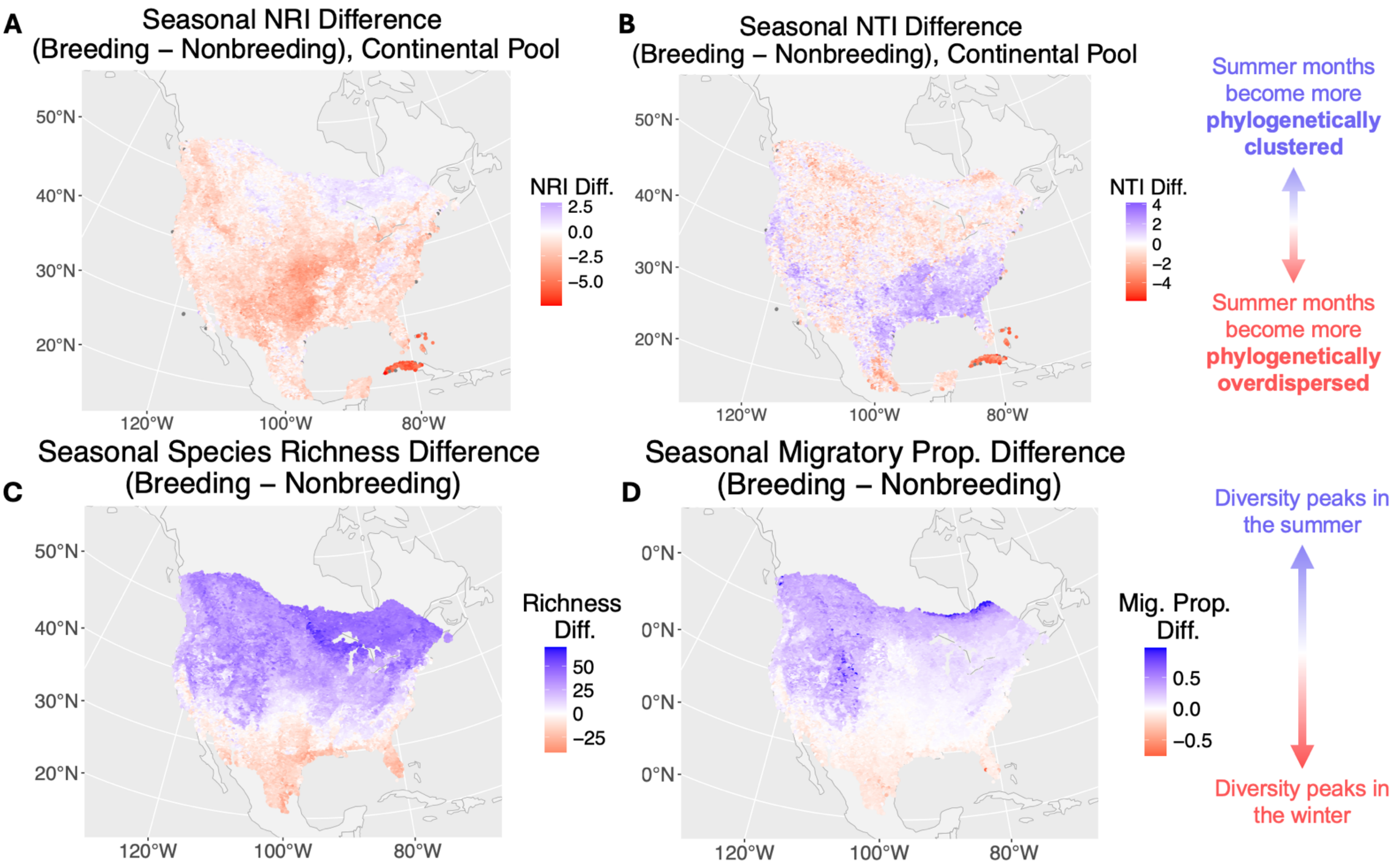


**Supplemental Figure S4:** Maps representing community seasonal differences in net relatedness index (NRI; A); nearest taxon index (NTI; B); species richness (C); and migratory proportion (D), using the continental species pool. Seasonal difference values represent breeding season values (using data from the last week of June) subtracted by non-breeding season values (first week of January). For both indices, blue values represent greater phylogenetic clustering during summer months as compared to winter months; red values depict greater phylogenetic overdispersion during summer. Blue and red values also represent higher species richness (C) and migratory proportion (D) during the breeding and non-breeding seasons, respectively.

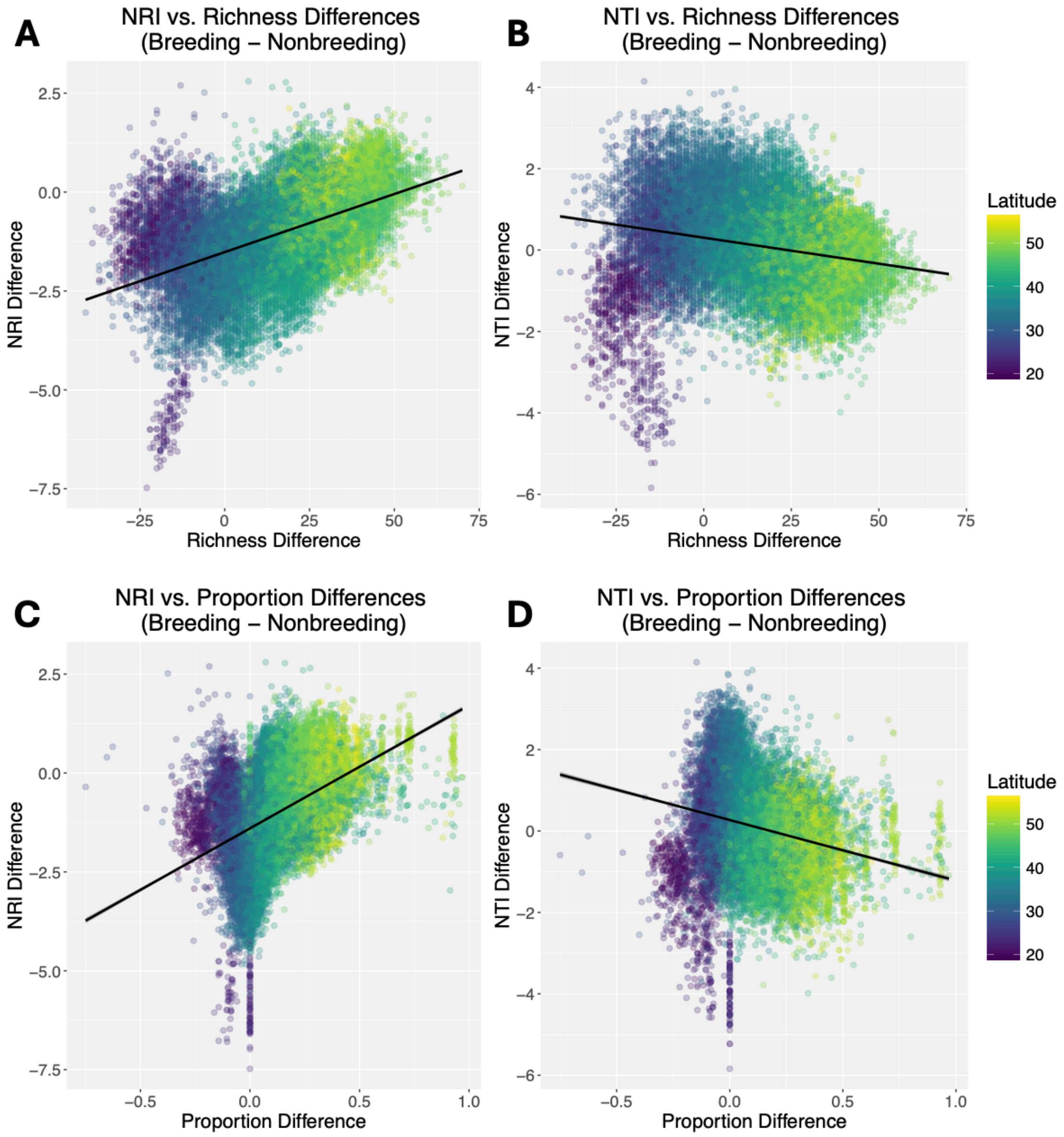


**Supplemental Figure S5:** Scatterplots depicting relationships between seasonal differences in community dispersion indices, and seasonal differences in species richness (A, B) and migratory proportion (C, D). Phylogenetic dispersion-community composition relationships are compared using the continental species pool. The left column (A, C) represents NRI relationships, while the right column (B, D) represents NTI relationships. Seasonal difference values represent breeding season values (using data from the last week of June) subtracted by non-breeding season values (first week of January). Spatial GLMMs predict that seasonal increases in richness and migratory proportion are associated with a seasonal increase in NRI (greater clustering) and a seasonal decrease in NTI (greater overdispersion). Standard errors are very low such that error bars around the trendlines are not visible.

| ΔNTI | Term | Estimate | SE | 95% CI |
|---|---|---|---|---|
| Richness | Intercept | 0.322 | 0.00905 | **[0.304, 0.339]** |
| | Richness | -0.0111 | 0.000412 | **[-0.0119, -0.0103]** |
| Proportion | Intercept | 0.284 | 0.00853 | **[0.268, 0.301]** |
| | Proportion | -1.22 | 0.0488 | **[-1.32, -1.13]** |
| Global | Intercept | 0.431 | 0.00965 | **[0.412, 0.450]** |
| | Richness | -0.00863 | 0.000579 | **[-0.00977, -0.00749]** |
| | Proportion | 1.10 | 0.0906 | **[0.921, 1.28]** |
| | Richness x Proportion | -0.0744 | 0.00262 | **[-0.0795, -0.0693]** |

| ΔNRI | Term | Estimate | SE | 95% CI |
|---|---|---|---|---|
| Richness | Intercept | -1.53 | 0.00826 | **[-1.55, -1.52]** |
| | Richness | 0.0214 | 0.000377 | **[0.0206, 0.0221]** |
| Proportion | Intercept | -1.46 | 0.00788 | **[-1.47, -1.44]** |
| | Proportion | 2.27 | 0.0453 | **[2.18, 2.36]** |
| Global | Intercept | -1.64 | 0.00877 | **[-1.66, -1.62]** |
| | Richness | 0.0167 | 0.000526 | **[0.0157, 0.0178]** |
| | Proportion | -0.678 | 0.0823 | **[-0.839, -0.516]** |
| | Richness x Proportion | 0.0714 | 0.00237 | **[0.0668, 0.0761]** |

**Supplemental Table S1:** Results of spatial generalized linear mixed models quantifying drivers of variation in seasonal turnover in NTI and NRI, with predictor variables of seasonal turnover in species richness and migratory proportion (along with their interaction). These results are derived from the continental species pool. Parameters whose 95% confidence intervals do not overlap with zero are bolded.

| **Predictor** | Absolute value of ΔNRI | Absolute value of ΔNTI |
|---|---|---|
| BIO4 (Temperature seasonality) | **0.000354 (0.000305-0.000403)** | 0.000033 (-0.000014 – 0.000081) |
| BIO7 (Annual temperature range) | **0.0161 (0.0143-0.0178)** | **-0.00710 (-0.00884 - -0.00536)** |

**Supplemental Table S2:** Effects of bioclimatic indices (temperature seasonality and annual temperature range) on the absolute value of ΔNRI and ΔNTI, using spatial generalized linear mixed models. These results are derived from the ecoregional species pool. Cell values represent predictor variable coefficients, with 95% confidence intervals in parentheses. Significant predictors ($p < 0.05$) are bolded.
